# A Novel Computational Framework to Investigate the Biomechanics of Burrowing

**DOI:** 10.64898/2026.10.10.758257

**Authors:** Jingrui Hu, Anastasiia Maliuk, Ce Liang, Sina Najjari, Sara-Adela Abad Guaman, Liwei Guo, Dominique Adriaens, Tiana Kohlsdorf, Henrik Birkedal, Anthony Herrel, Susan E. Evans, Mehran Moazen

**Affiliations:** Department of Mechanical Engineering, University College London, London, UK; Faculty of Agricultural Sciences and Renewable Natural Resources, Universidad Nacional de Loja, Loja, Ecuador; Evolutionary Morphology of Vertebrates, Department of Biology, Ghent University, Ghent, Belgium; Department of Biology, FFCLRP, University of São Paulo, São Paulo, Brazil; Department of Chemistry, Aarhus University, Aarhus, Denmark; Mécanismes Adaptatifs et Évolution, UMR 7179, Muséum national d’Histoire naturelle, CNRS, Paris, France; Functional Morphology Laboratory, Department of Biology, University of Antwerp, Antwerp, Belgium; Naturhistorisches Museum Bern, Bern, Switzerland; Department of Cell and Developmental Biology, University College London, London, UK

**Keywords:** Biomechanics, craniofacial system, amphisbaenians, head-first burrowing, discrete element method, finite element analysis

## Abstract

Head-first burrowing likely imposes strong mechanical demands on the skull of fossorial vertebrates, but how cranial tissues respond to these loads remains obscure. Here, we developed a computational framework integrating the discrete element method (DEM) with finite element analysis (FEA) to investigate cranial biomechanics during burrowing. We used *Blanus cinereus*, a round-headed amphisbaenian, as a model to establish DEM predictions that were validated and calibrated against *in vitro* experiments across three substrates and compared with *in vivo* push forces. DEM-calibrated particle contact forces were then transferred to an anatomically detailed finite element model to quantify cranial loading. The analyses suggested that soft tissues accommodated substantial deformation, while the skull transmitted burrowing loads from the premaxilla towards the occipital region. Cranial sutures further redistributed stresses and strains within the skull. This validated DEM-FEA framework links particle-scale substrate interactions with internal cranial biomechanics and provides a quantitative approach for investigating the mechanics of head-first burrowing across fossorial vertebrates.

## 1. Introduction

Burrowing is a fundamental locomotor strategy for many fossorial organisms, enabling underground movement that provides access to shelter, foraging opportunities, and protection from predators and environmental stress ^1^. Amphisbaenians are fossorial vertebrates that use the skull as the primary structure for burrowing ^2^, and simultaneously exhibit substantial cranial diversity, including four principal head morphologies suggested to be associated with different burrowing behaviours ^3–9^. Several of these cranial morphotypes have evolved independently across amphisbaenian lineages, suggesting convergent responses to the functional demands of head-first burrowing ^6,10^. These observations suggest that skull-substrate interactions are likely to be central to understanding such convergence ^11,12^. However, the cranial biomechanics underlying these different burrowing behaviours remains poorly understood.

The main challenge of such investigations arises from characterising the dynamic mechanical response of the cranial system to external forces while preserving the complexity of skull-substrate interactions. For example, *in vivo* experiments provide valuable measurements of external burrowing forces and motion, but cannot directly quantify the force distribution onto the skull ^11,13^. *In silico* experiments can be effective for assessing structural mechanical behaviour, but are limited in their ability to reproduce the real, particle-scale interactions between the head and the surrounding substrate. Together, these limitations motivate a modelling approach that can capture the discrete nature of the burrowing medium.

The discrete element method (DEM) is a numerical technique that provides a potential solution to this problem by explicitly modelling interactions among individual substrate particles and between particles and biological structures ^14^. When coupled with finite element analysis (FEA), DEM-derived contact forces can be used to quantify how realistic substrate interactions generate internal stresses and strains within biological tissues. It has been widely applied in both basic research and industry to solve problems involving granular and particulate materials, including mining ^15^, agriculture ^16^, and civil engineering ^17^. However, to date, this method has not been used to integrate substrate contact forces with internal cranial biomechanics. As such, rigorous validation is required before it can be applied to biomechanical questions related to head-first burrowing.

In this study, we used *Blanus cinereus* (Blanidae, Amphisbaenia) as a representative of a generalised round-headed head-first burrower, considered the ancestral cranial condition in amphisbaenians ^18^, to establish and validate a biomechanical framework for head-first burrowing. We first validated DEM-predicted global and local forces against *in vitro* experiments. The *in vitro* set up involved fixing a scaled-up 3D printed skull to a material testing machine and driving the skull model into three different media, PMMA (Poly (methyl methacrylate)) spheres, sand, and soil. We then developed a more detailed anatomical model including various soft tissues within the head (skin, muscles, and sutures) and compared model predictions with *in vivo* burrowing forces. The particle-scale contact forces predicted by DEM were then transferred directly to an anatomically detailed FE model, enabling the resulting cranial stress, strain and load transfer to be quantified under realistic skull-substrate interactions. By using this framework, we identified the distinct mechanical roles of the cranial soft tissues, sutures, and bones during burrowing. More importantly, our work established a robust approach for integrating realistic granular interactions with internal tissue biomechanics, and it can be readily transferred to other fossorial locomotion studies.

## 2. Results

### 2.1 Experimental Validation of the Discrete Element Method Framework

The resultant forces measured by the load cell (Instron) were first compared with the reaction forces from the DEM simulations (Figure 1a). For the PMMA and sand substrates, the overall trend and peak force magnitudes showed good agreement throughout the burrowing process, although the DEM force exhibited greater fluctuations than the experimental measurements (Figure 1b and 1c). Detailed *R*^2^, Pearson *r*, and RMSE values are provided in Supplementary Table S1. For soil, the DEM parameters were calibrated against the *in vitro* measurements, with the best-fitting parameter set listed in Supplementary Table S2. The DEM simulations slightly overestimated the reaction forces during the intermediate stage of penetration (1 – 4 s) but underestimated them during the final stage (> 4.5 s), as shown in Figure 1d. Nevertheless, the overall agreement between experimental and simulated force responses was high (*R²* = 0.93, Pearson *r* = 0.97, RMSE = 4.27 N). These parameters were subsequently used for the DEM simulations designed to mimic the *in vivo* experiments.

**Figure 1.**
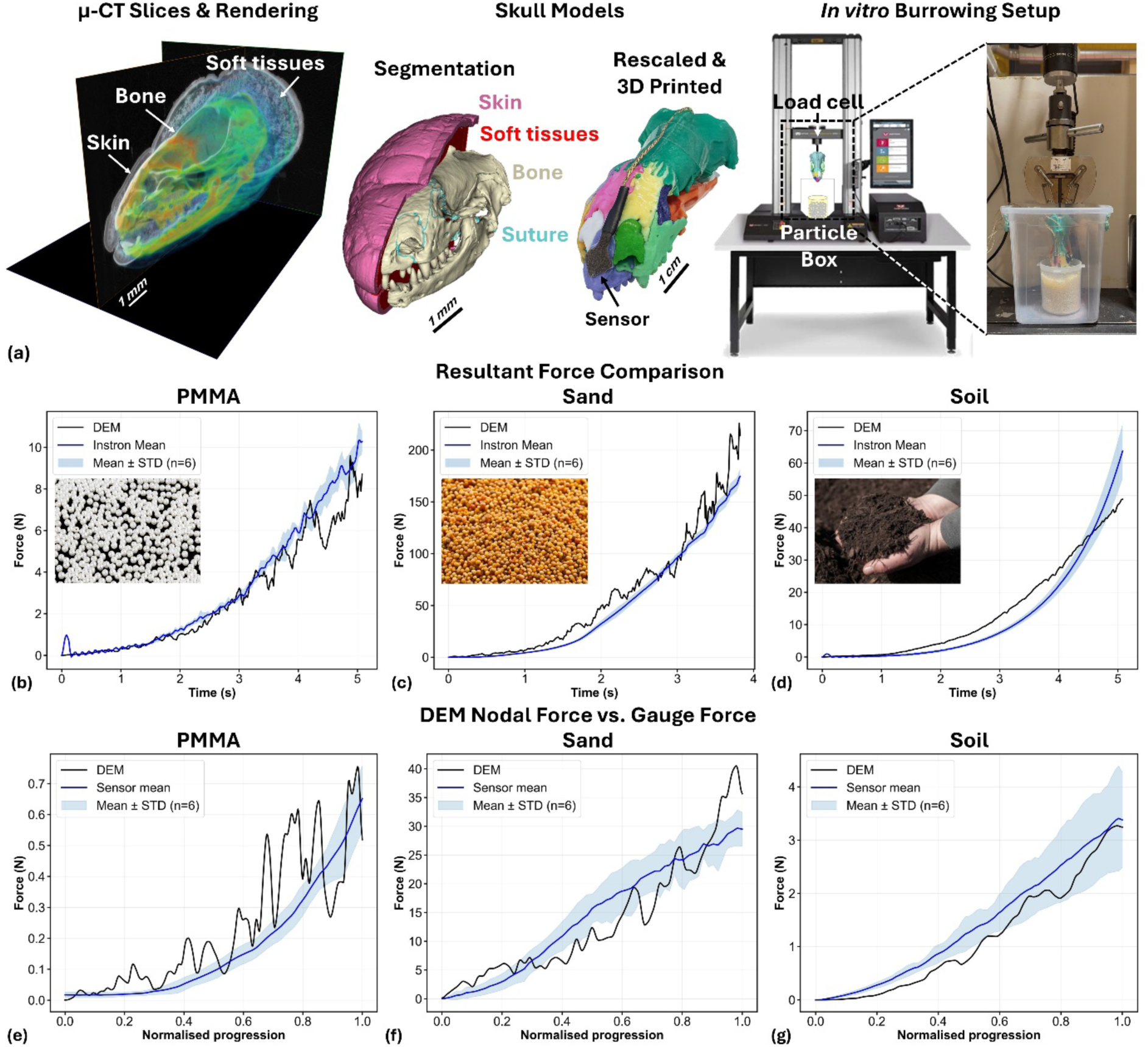
(a) Overview of the experimental and computational workflow. The µCT images of *Blanus cinereus* were segmented, rescaled, and used to generate a 3D-printed model. The printed model was then mounted on the Instron mechanical testing machine for in vitro burrowing experiments across three different substrates. (b – d) Comparison between the resultant reaction forces measured experimentally using the Instron load cell (blue) and those measured from the DEM simulations (black) for PMMA, sand, and soil substrates. (e – g) Comparison between the DEM-derived nodal forces and forces (black) measured by the local pressure sensors (blue). Solid lines represent mean experimental values, and shaded regions indicate standard deviation. The DEM model reproduced the overall force progress and magnitude observed experimentally across the tested substrates.

In terms of the local force, comparisons between the DEM-derived nodal forces and those measured by the sensor revealed generally consistent trends across all substrates (Figure 1e – g). For PMMA, the DEM nodal forces exhibited greater fluctuations than the *in vitro* measurements, although the overall trend was captured (*R*^2^ = 0.54, Pearson *r* = 0.87). A similar pattern was observed for sand, but with fewer fluctuations (*R^2^* = 0.81, Pearson *r* = 0.92). In contrast, the soil substrate exhibited substantially less fluctuation, and the DEM results closely matched the experimental data, yielding the highest level of agreement among the three substrates (*R^2^* = 0.92, Pearson *r* = 0.99).

### 2.2 Burrowing Force Generation in *Blanus cinereus*

Two *in vivo* tunnels of different lengths (20 and 200 mm) were used to measure the push forces generated by living *Blanus cinereus* (*B. cinereus*), which were then compared with the forces predicted by the DEM simulations. At the initial burrowing stage (20 mm tunnel), *B. cinereus* primarily generated forward-directed forces during burrowing with maximum forward forces reaching approximately 0.3 N. Horizontal forces were smaller, with peak values around 0.1 N, and the vertical forces were negligible throughout burrowing. The DEM simulations and short tunnel *in vivo* experiments exhibited comparable force magnitudes and similar temporal force patterns, except that the vertical force in Z direction was slightly over predicted (Figure 2c).

**Figure 2.**
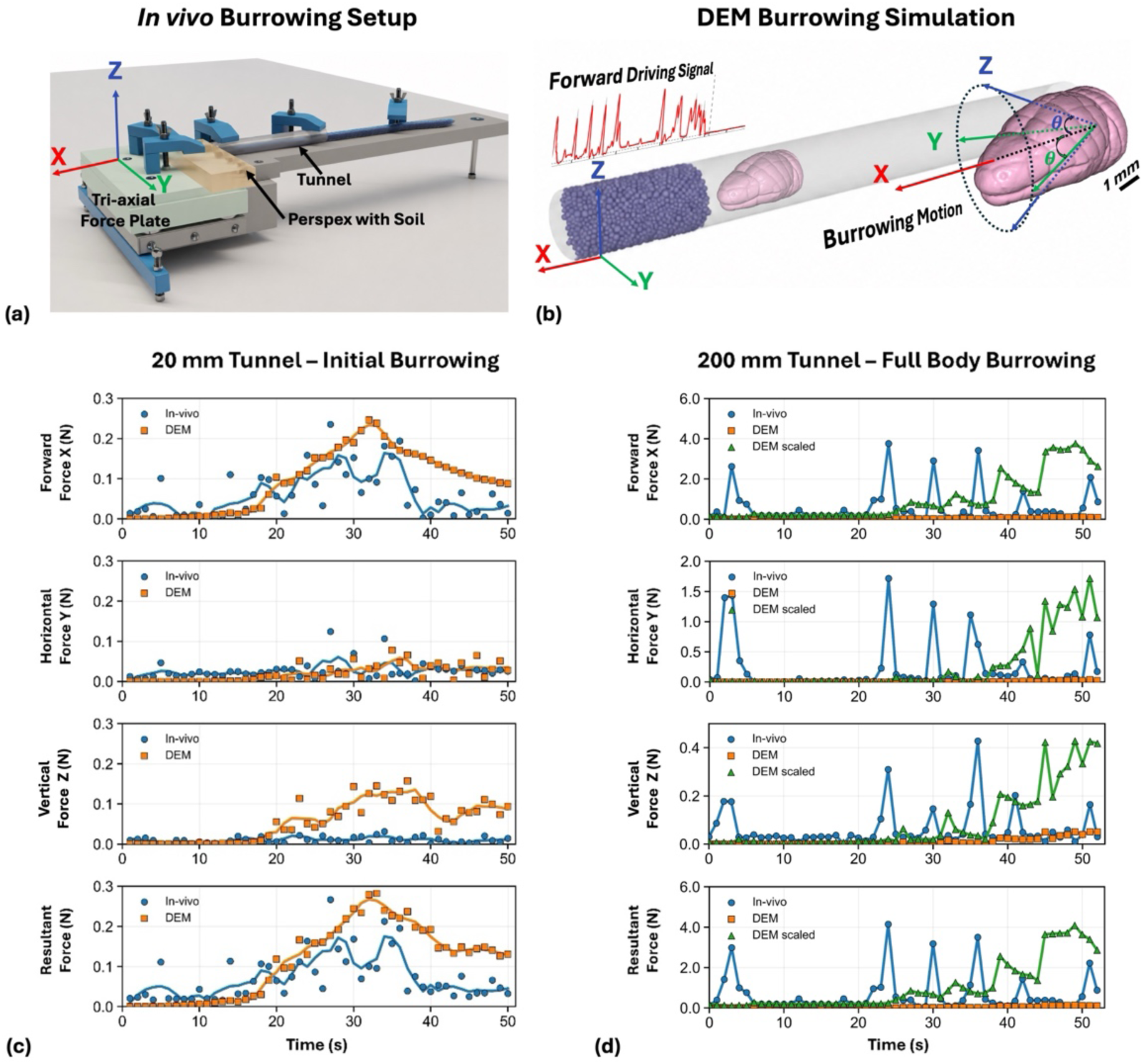
(a) Schematic of the *in vivo* burrowing force measurement setup. The individual was positioned within a transparent tunnel, and it pushed into soil contained within a Perspex block. The Perspex block was mounted on a tri-axial force plate to record forces in three directions, and the entire system was supported by a custom-built metal frame. (b) DEM simulation replicating the *in vivo* experimental conditions shown in (a). The prescribed burrowing kinematics comprised forward translation along the X-axis and rotations about the Y-and Z-axes, represented by two rotational degrees of freedom (|*θ*| ≤ 1°). *In vivo* measured forces were compared with DEM-predicted forces using (c) a short tunnel (20 mm), representing the initial stage of burrowing in which only the *Blanus cinereus* (*B. cinereus*) skull interacted with the substrate; and (d) a long tunnel (200 mm), in which the whole body could contribute to force generation, allowing measurement of the maximum forces generated by *B. cinereus*. From top to bottom, graphs show the forward (X), horizontal (Y), vertical (Z) forces, and resultant forces, respectively. Blue curves represent the *in vivo* measurements, orange curves represent the DEM simulation results, and green curves represent the scaled DEM simulation results.

During the full-body burrowing scenario (200 mm tunnel), the measured *in vivo* forces were greater than the DEM-predicted forces in all directions (Figure 2d). The discrepancy was most pronounced in the forward direction, where the peak force measured *in vivo* was 9.81 times greater than that of the DEM simulations. The horizontal force components in the *in vivo* test were 5.26 times greater than in the DEM simulation, while the vertical force showed considerably better alignment, with peak forces only 1.24 times higher than the corresponding DEM results. Using these peak-force ratios as scaling factors, the scaled DEM nodal forces reached peak magnitudes comparable to those of the *in vivo* measurements (Figure 2d, green vs. blue curves), although differences in the temporal force patterns remained.

### 2.3 Mechanical Roles of Cranial Tissues During Burrowing

The scaled DEM nodal forces were subsequently used to investigate the underlying cranial biomechanics. At the peak loading scenario, the skin and soft tissues underwent substantial deformation (Figure 3a and b), particularly compression, with averaged minimum principal strains of-0.12 and-0.11, respectively. Within the skull (Figure 3c, 3e and 3f), different bone compartments exhibited distinct patterns of principal strain. The premaxilla and nasal bones were predominantly under compression, while the frontal bones experienced mainly tensile strain. In the parietal bone, the lateral regions were predominantly under compression, whereas the dorsal ridge experienced tensile strain. The occipital bone showed a more complex strain pattern, with tensile strain along the lateral regions and both high compressive and tensile strains in the ventral region (Figure 3c and Figure S1). Notably, the cranial sutures experienced substantial deformation in both compression and tension. Among the sutures, the parietal-prootic sutures exhibited the greatest deformation, reaching a peak minimum and maximum principal strain of approximately-0.54 and 0.4, respectively. Followed by the frontoparietal suture, which had an average minimum and maximum principal strain of-0.042 and 0.056, respectively.

**Figure 3.**
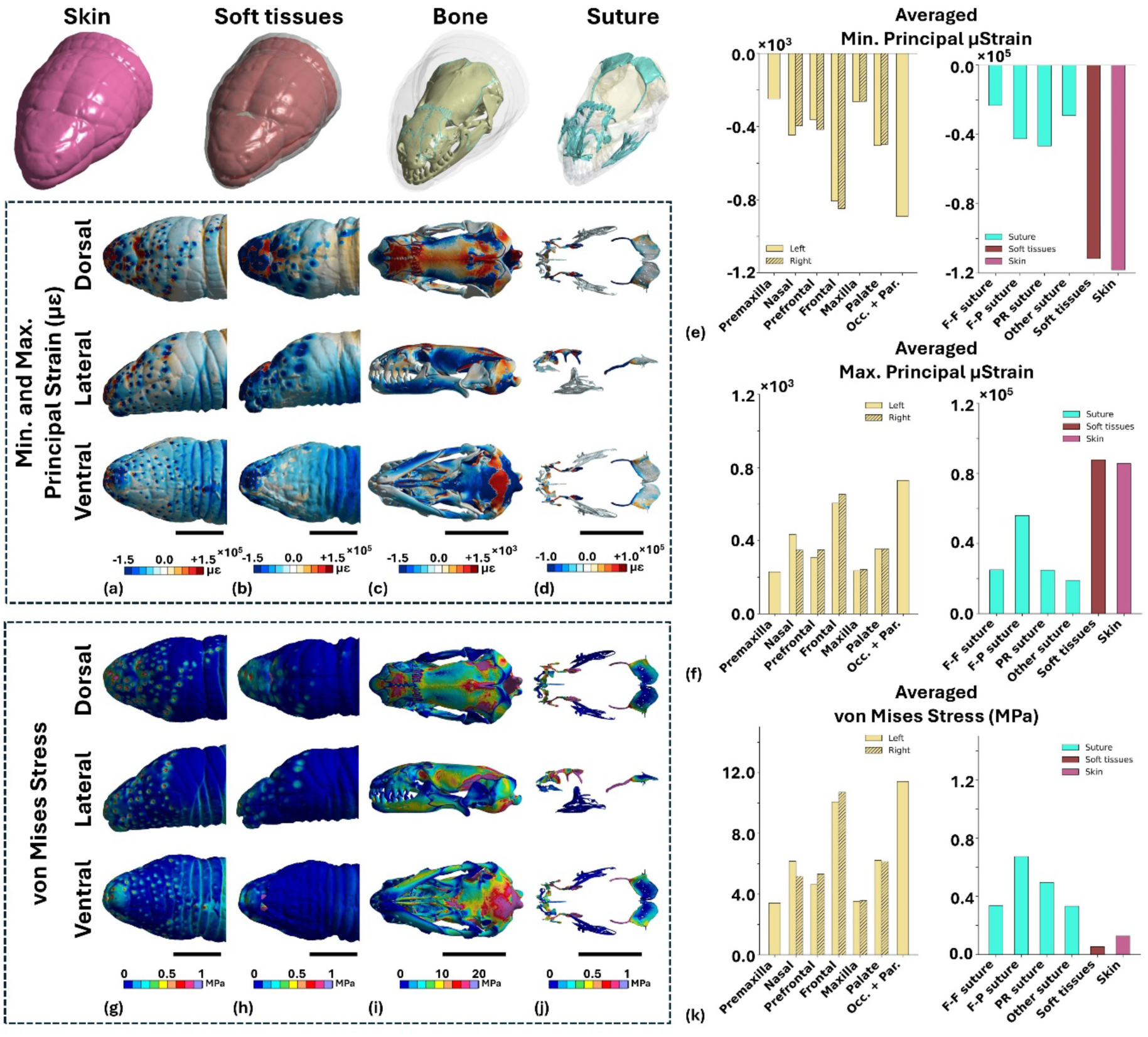
Distribution of minimum and maximum principal strain and von Mises stress in the skin, soft tissues, skull, and sutures of the *Blanus cinereus* skull model. (a – d) Minimum and maximum principal strain distributions shown in dorsal, lateral, and ventral views, with negative and positive values representing compressive and tensile strains, respectively. Corresponding average values are shown in (e) and (f), respectively. (g – j) Von Mises stress distributions shown in dorsal, lateral, and ventral views, with the corresponding average values shown in (k). Pr. suture – premaxilla-nasal suture; F-F. suture – interfrontal suture; F-P suture – Frontoparietal suture. Scale bar = 3 mm.

In terms of the von Mises stress distribution (Figure 3g – 3k), the maximum stress was observed at the occipital condyle, where the model was constrained. Excluding the boundary region, the frontal bones experienced the highest stress, with average values of 10.71 MPa on the right side and 10.07 MPa on the left. Interestingly, despite being the first cranial bone to directly interact with the substrate during burrowing, the premaxilla experienced relatively low stress, with an average value of only 3.41 MPa. The stress was particularly low at the tip of the premaxilla. Localised stress concentrations were also observed at the parietal-prootic bone-suture interfaces.

To better understand the function of cranial sutures during burrowing, the suture regions were virtually fused by assigning them the same material properties as bone. As shown in Figure 4, fused suture regions increased the stress and strain distributed in the corresponding bones. In particular, fusion of the frontoparietal suture increased the average stress by 27% in the left frontal bone and 52% in the right frontal bone, while the peak stress more than doubled (Figure 4a and 4c). Strain concentrations were also observed at the frontal-parietal interface following fusion of the frontoparietal suture (Figure 4b, 4d and 4e). Fusion of all cranial sutures established more extensive stress and strain concentrations, especially in the prefrontal bones (Figure 4b, 4d and 4e). The detailed values were provided in Supplementary Figure S2 and S3.

**Figure 4.**
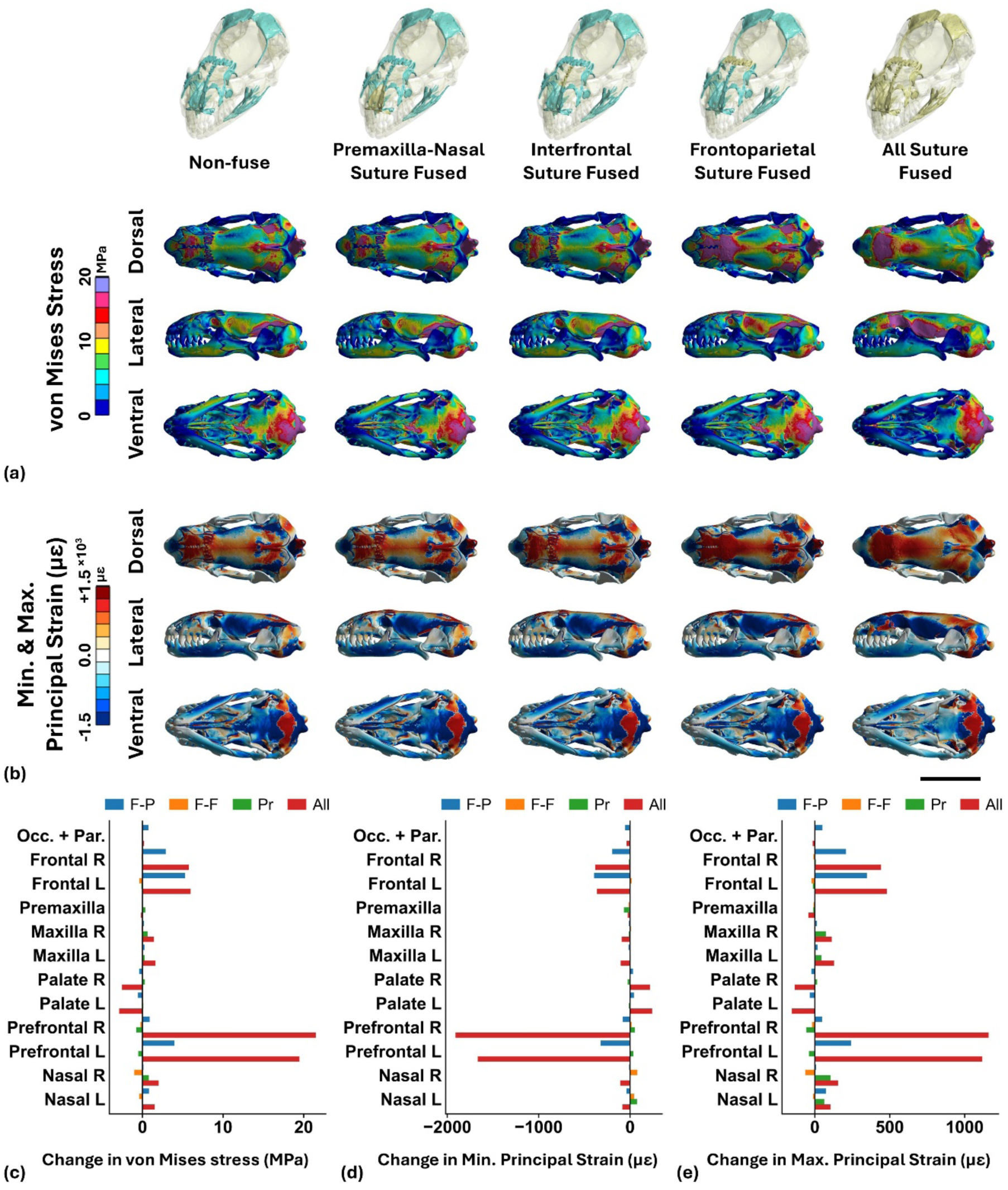
Distribution of von Mises stress (a) and minimum and maximum principal strains (b) in the skull and sutures of the *Blanus cinereus* skull model under different suture-fusion conditions. The skull is shown in dorsal, lateral, and ventral views. (c – e) Corresponding changes in average von Mises stress and minimum and maximum principal strains under different suture-fusion conditions. Pr. suture – premaxilla-nasal suture; F-F. suture – interfrontal suture; F-P suture – Frontoparietal suture. Scale bar = 3 mm.

## 3. Discussion

This study presents a novel framework that integrates, for the first time, the discrete element method (DEM) with finite element analysis (FEA) to investigate the skull biomechanics of head-first burrowing. The DEM model was first validated with *in vitro* experiments by comparing both local nodal forces and the resultant reaction forces. It was then calibrated using *in vivo* measurements obtained the initial penetration and the whole-body contributions to force generation. The calibrated nodal forces were subsequently applied as loading conditions in the finite element model, enabling a more realistic evaluation of skull biomechanics during burrowing. This framework has a major contribution of providing new insights for investigating biomechanical adaptations to fossorial locomotion, and can be readily extended to other burrowing animals.

Validation against the *in vitro* experiments demonstrated that the DEM can reliably reproduce both the global reaction force and local contact forces. Experimental measurements were consistent with DEM predictions across three tested substrates, including different particle morphologies and contact behaviours (Figure 1). Importantly, agreement between the measured and DEM-predicted local forces provides support for the model’s representation of contact loading at the premaxillary region (Figure 1e – g). This is particularly important for subsequent FEA, where the predicted mechanical response strongly depends on the accuracy of local loading conditions. The larger fluctuations observed for the DEM predicted PMMA nodal forces (Figure 1e) likely result from its relatively large and smooth particles. The relatively small number of particles simultaneously contacting the skull model surface made the local contact forces highly sensitive to individual particle rearrangements, rolling, and the continual progress of the contact network ^14,19–21^. This interpretation is further supported by the particle size and friction sensitivity analyses as shown in Supplementary Figure S4 and S6.

The finer sand and soil particles generated a denser and more continuous contact network, resulting in a smoother force pattern. Although the particle size of sand used for the DEM simulation was approximately ten times larger than that used experimentally, the simulated force curve reproduced the overall temporal pattern and force magnitude, despite showing slightly larger fluctuations (Figure 1c and 1f). Particle upscaling may alter the number and organisation of contacts at the head-substrate interface. Therefore, the similarity between the simulated and experimental force responses supports the DEM model at the macroscopic force level, while the microscale contact fluctuations should not be interpreted as directly representative of the experiment ^22^. For more cohesive soil, discrepancies between the experimental and DEM reaction force profiles remained (Figure 1d and 1g). Such divergence probably resides on the fact that the soil used in the *in vitro* experiments comprising not only mineral particles but also organic fibres, mulch, and manure, whereas the DEM model represented the material using uniform spherical particles with simplified contact mechanics. Consequently, the simulation did not effectively capture the bulk responses of the constituents, which contributed to the higher reaction forces observed during the initial penetration stage and the lower forces at later stages in the simulation ^14,19,21,23^.

For *in vivo* tests, during the initial stage of burrowing, the peak forces predicted by the DEM were comparable in magnitude to those measured in live specimens (Figure 2c), although the temporal force pattern did not precisely overlap. This difference is expected because the DEM calculates substrate reaction forces resulting from the prescribed head motion ^24,25^, whereas the *in vivo* measurements reflect forces actively generated by the *Blanus cinereus* (*B. cinereus*) and transmitted to the substrate through the head ^11,13^. In the short tunnel, the neck was the primary structure generating push force, and therefore the peak forces were similar in both DEM and *in vivo*. However, when *B. cinereus* were pushing along a long tunnel, the axial musculature and body-wall interactions contribute substantially to locomotion, enabling the animal to generate considerably greater forces ^11,13^. In this scenario, the *in vivo* measurements represent the maximum pushing forces that *B. cinereus* can produce during full-body burrowing. Consequently, the DEM-predicted force magnitudes were much lower than those measured *in vivo*. Nevertheless, because the DEM provides the spatial distribution of contact forces acting on the FE model, scaling these nodal forces to the experimentally measured peak load still enabled biologically realistic loading conditions for subsequent finite element analyses, by projecting the temporal DEM nodal forces onto the finite element (FE) model.

While experimental studies have extensively quantified burrowing performance and external forces, relatively few studies have used computational modelling to investigate internal biomechanical responses, partly because realistic soil-body interactions are difficult to represent. Previous FEA studies have therefore generally relied on simplified or prescribed external loading conditions, which may not fully capture the spatial and temporal complexity of substrate-induced loading during burrowing ^26,27^. Our results suggest that the *B. cinereus* cranial system functions as a composite structure, in which the soft tissues, including the skin, accommodate the majority of the deformation. Previous studies have emphasised the importance of the skin ^28,29^ and muscles ^2,28,30^ in facilitating locomotion and burrowing, but their contribution to cranial load has not been quantified. Our findings indicate that, beyond providing locomotor flexibility and force generation, these soft tissues may also function as the primary mechanical buffer, accommodating deformation and redistributing loading around the underlying skull. In contrast, the skull bone is mechanically rigid during burrowing, and transfers load efficiently from the premaxilla (tip of the snout) towards the occipital condyle. This interpretation is consistent with the consolidated and less kinetic skull morphology of amphisbaenians described in previous studies ^11,28,30^. These behaviours are also consistent with established principles of tissue biomechanics, whereby compliant soft tissues undergo greater deformation under loading, whereas hard tissues primarily support and transmit mechanical loads ^31,32^. Although extensive previous studies have also highlighted the functional significance of cranial morphology for head-first burrowing ^5,6,9,10,33–35^, they have primarily inferred mechanical function from anatomical observations. Our study provides a more realistic description of cranial load transfer during head-first burrowing, and we expect that this framework may help to explain the mechanical basis of specialised cranial architecture in the future.

The tip of the snout experienced relatively low stresses despite being the first region to contact the substrate. This may partly reflect the greater amount of soft tissue surrounding the snout, which accommodates deformation and buffers the underlying bone. The low stress may also be related to the conical geometry of the premaxillary bone, as more pointed head shapes have been shown to require lower forces for substrate penetration ^8^. Burrowing loads are subsequently transmitted posteriorly through the frontal bones towards the parietal and occipital regions, where higher stress concentrations were observed (Figure 3). As *B. cinereus* is a round-headed forward burrower, these results provide biomechanical insights into the load transfer path and support for the hypothesis that the round characteristic head shape represents an adaptation for this burrowing strategy ^6,9,11,35,36^. Future studies will extend this framework to species with different cranial morphologies and substrate types to investigate how variation in cranial shape influences biomechanical performance during burrowing.

Notably, the cranial sutures play an important role in amphisbaenian burrowing by significantly reducing stress concentration on skull bones, as demonstrated in Figure 3 and 4. Pronounced stress concentrations were established in the frontal, parietal and prefrontal bones after digital fusion of the sutures, indicating that the sutures help to redistribute mechanical loads during burrowing (Figure 4a and 4c). Previous studies of amphisbaenians have primarily described cranial sutures from anatomical and morphological perspectives, whereas their mechanical function remained unexplored ^30^. Our results therefore provide further biomechanical evidence that cranial sutures contribute to the burrowing-induced loads by providing sufficient flexibility to dissipate mechanical stress while maintaining the overall stiffness of the skull, which is consistent with experimental and computational studies in other animals ^37–40^. It is worth noticing that the fusion of the more anterior sutures around the premaxilla and frontal region (interfrontal suture) produced comparatively small changes in bone stress distribution, despite their proximity to the initial substrate contact. In contrast, fusion of the frontoparietal suture produced a much greater mechanical response. This suggests that the mechanical importance of individual sutures is defined not simply by their proximity to the applied load, but by their position within the cranial load-transfer pathway, which is closely related to their burrowing motion. The frontoparietal interface may therefore represent a particularly important compliant junction through which anteriorly generated burrowing loads are redistributed to the posterior skull.

We recognise limitations of this study, which do not minimise the clear contribution of the validated DEM-FEA framework to the field of biomechanics in fossorial species. Firstly, the DEM forces are largely determined by model motion, while the *in vivo* experiments measured the maximum push forces generated by amphisbaenians, rather than forces produced during natural burrowing. Consequently, the derived motion represents only part of the loading experienced by the skull during burrowing. Nevertheless, the simulations employed rescaled loading conditions that reasonably captured the biomechanical response of the skull. Future work will incorporate X-ray videography (e.g., XROMM ^41^ or biplanar fluoroscopy ^42,43^) to accurately quantify burrowing kinematics and integrate muscle-generated forces, enabling a more comprehensive analysis of cranial biomechanics. Secondly, only one specimen was scanned and modelled, and therefore potential biological variation associated with age, sex, and individual morphology was not considered. Future studies incorporating several specimens will allow assessing the influence of such variation on cranial biomechanics. Thirdly, several DEM contact parameters, particularly those describing interactions between the substrate and the skull model, could not be directly measured or obtained from the literature. Although the adopted parameter set successfully reproduced the experimentally measured force responses, alternative parameter combinations may produce similar macroscopic behaviour. Future studies focusing specifically on substrate characterisation may employ dedicated parameter calibration experiments to further constrain these contact parameters ^44^.

In conclusion, this study presents the first DEM-FEA framework for investigating the biomechanics of head-first burrowing in amphisbaenians. Validation against both *in vitro* and *in vivo* experiments demonstrated that the model can realistically reproduce skull-substrate interactions and generate biologically relevant loading conditions. The biomechanical analyses revealed that the *B. cinereus* cranial system functions as a composite structure, in which the skin and soft tissues act as a mechanical buffer, the cranial sutures distribute mechanical loads, and the rigid skull efficiently transmits loads from the premaxilla to the occipital region. Together, these findings provide a quantitative description of cranial load transfer during simulated head-first burrowing and demonstrate the contrasting mechanical roles of bone, sutures, and soft tissues within the round-headed *B. cinereus* head. The framework provides a basis for future comparative studies examining how variation in cranial morphology influences biomechanical performance across amphisbaenians with different head shapes and burrowing strategies.

## 4. Methodology

### 4.1 Sample Preparation and Model Development

One alcohol preserved amphisbaenian specimen of – *Blanus cinereus* (*B. cinereus*) from the research collection of Anthony Herrel was scanned using micro-computed tomography (Figure 1a – left, µCT, Nikon XTH 225 ST scanner, Nikon Metrology Ltd., Japan). Scanning was performed at a voxel size of 6 μm, with an X-ray beam energy of 65 kV and a power setting of 7 W. The reconstructed image dataset was used for subsequent processing using Simpleware (X, Synopsys Inc., CA, US) to segment the skull, skin, suture and soft tissue (Figure 1a – middle). The segmented skull (without mandibles) was converted into a stereolithography (STL) file, then scaled to 15 times its original size, and 3D printed using a PolyJet 3D printer (J835, Stratasys Ltd., MN, US). The skull was printed in rigid photopolymer materials (VeroVivid, Stratasys Ltd., MN, US), with different anatomical structures represented by distinct colours (Figure 1a – middle).

The preserved specimen was subsequently stained with phosphomolybdic acid (PMA) and rescanned using the same µCT setup to facilitate segmentation of the skin and other soft tissues. The segmented skull model obtained from the unstained specimen, which provided better contrast for the skull and cranial sutures, was rigidly registered to the stained scan and integrated with the segmented skin and soft tissues to reconstruct the complete head model (Figure 1a-left and middle).

### 4.2 *In vitro* Experimental Setup

For the *in vitro* experiment, the 3D-printed model was mounted on an Instron mechanical testing machine (Model 5985, Instron, MA, US) and driven vertically into a particle-filled box from top to bottom at a constant speed of 0.01 m/s for 5 s, as shown in Figure 1a – right. The compression process was repeated six times, with the particles reset to their initial configuration prior to each run. The total resultant force was recorded using a 500 N load cell. Local particle forces were measured using a tactile pressure sensor (ConTacts C500, Pressure Profile Systems, Inc., CA, US) attached to the surface of the 3D-printed model (Figure 1a-middle). The sensor had an active measurement area of 10 × 10 mm² and was positioned at the anterior tip of the premaxilla, where compressive loading was expected to dominate. The sensor was powered at 5 V, and the output was acquired using an ESP32 microcontroller (FNK0060, Freenove, Shenzhen, China) with its integrated 12-bit analogue-to-digital converter and transmitted to a computer for recording using a custom Python script developed in Visual Studio Code (Microsoft Corporation, WA, US). Prior to the measurements, the sensors were calibrated using reference weights of 10, 20, 50, and 100 g.

Three different materials were selected for validation: 1) Poly (methyl methacrylate) (PMMA) beads, representing the simplest homogeneous material with negligible rolling resistance; 2) desert sand, representing a heterogeneous material with mixed particle shapes and sizes; and 3) commercial potting soil, representing a more realistic substrate with adhesive interactions.

### 4.3 *In vivo* Experimental Setup

*In vivo* burrowing forces were measured using a custom-designed piezoelectric (Kistler) force platform, as shown in Figure 2a. The detailed description of the setup was published ^11,13^. In brief, a live *B. cinereus* individual was placed in a tunnel with a diameter equal to the maximum body width (6 mm). Two tunnel lengths were used. The 20 mm tunnel represented the initial stage of burrowing, during which only the head interacted with the substrate. The 200 mm tunnel represented full-body burrowing, during which the entire body entered the tunnel and was used to measure the maximum pushing forces generated by *B. cinereus*. The tunnel was positioned in front of a Perspex block containing a hole with the same diameter as the tunnel and filled with a 5 mm layer of soil. The Perspex block was mounted on a force plate to measure forces when *B. cinereus* pushed into the soil.

The measured burrowing forces were then used to derive the burrowing motion used for discrete element method (DEM) simulation. The force signals were low-pass filtered (10 Hz cutoff), and the forward force component was converted into a normalised forward velocity profile according to Eq. 1,

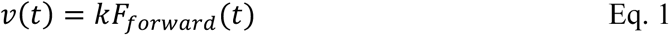

where *v*(*t*) was the prescribed forward velocity, *F_for_*_w*ard*_(*t*) was the forward force, and *k* (mm·N^-1^·s^-1^) represented a scaling factor calibrated to ensure that the total forward displacement matched the burrowing distance measured experimentally (5 mm). The resulting velocity profile was applied directly as the translational motion in the DEM simulations.

The lateral and vertical force components were normalised and mapped to left-right and up-down head rotations through a first-order dynamic response model, with rotation amplitudes limited to ± 1° by assumption (Eq. 2). Since direct measurements of head kinematics were not available, the rotational motion should be regarded as an approximation rather than an exact reconstruction of the animal’s movement,

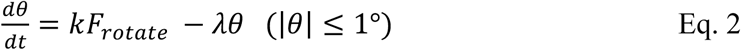

where *F_rotate_* was the normalised force component, *θ* was the rotation angle, and *λ* was the decay constant (6 s^-1^). The rotational scaling factor was defined as *k* = *λθ_max_*. The resulting translational and rotational motions were converted to velocity and angular velocity inputs for the DEM model.

### 4.4 Discrete Element Method Simulation

The discrete element method (DEM) simulation was run using Rocky (2025R2, ANSYS Inc., PA, US). The material properties of each particle type were summarised in Table 1. The bulk density was determined by measuring the mass of particles occupying a known volume. The bulk solid fraction was then calculated as the ratio of bulk density to particle density, providing a measure of the proportion of the bulk volume occupied by solid particles.

**Table 1.** The material properties of validation particles in DEM simulations.

| Property | DEM |  |  |  | FEA |  |  |  |
| --- | --- | --- | --- | --- | --- | --- | --- | --- |
|  | PMMA | Sand | Soil | Skull (Printed) | Skull (Bone) | Suture | Skin | Soft Tissue |
| **Bulk Density (kg/m<sup>3</sup>)** | 570 | 1,670 | 600 | — | — | — | — | — |
| **Bulk Solid Fraction** | 0.50 | 0.63 | 0.60 | — | — | — | — | — |
| **Density (kg/m<sup>3</sup>)** | — | — | — | 1,200 | 1,900 | 1,050 | 1,100 | 1,000 |
| **Young's Modulus (MPa)** | 3,000 | 70,000 | 50 | 2,000 | 10,000 | 10 | 1 | 0.45 |
| **Poisson's Ratio (-)** | 0.37 | 0.30 | 0.30 | 0.30 | 0.30 | 0.30 | 0.40 | 0.45 |
| **Particle Diameter (mm)** | 4.0 | 2.0 | 0.5 | — | — | — | — | — |
| **Reference** | 45 | 45,46 | 47 | 48 | 49 | 49 | 50 | 51 |

The PMMA particles were modelled using a linear spring-based contact model, comprising a hysteretic linear spring normal force law and a linear spring coulomb tangential force limit. Adhesion and rolling resistance were not included, reflecting the smooth and near-spherical morphology. This contact formulation has been widely applied for cohesionless granular materials ^24,52^. The sand particles were modelled using the same contact laws but with linear spring rolling resistance at 0.2 to mimic the slight effects of shape and roughness. As the physical sand particles were approximately 0.05 – 0.2 mm in size, making direct representation computationally prohibitive, a representative particle size of 2 mm was selected based on sensitivity tests to balance computational feasibility and model stability (Supplementary Figure S4 and Table S3). The detailed particle-particle parameters were recorded in Supplementary Table S2.

For the soil, particle interactions were modelled using a nonlinear model with a hertzian spring-dashpot normal contact law and a linear spring coulomb tangential friction law. The Johnson-Kendall-Roberts (JKR) adhesive model together with a linear spring rolling limit rolling resistance models were incorporated to account for interparticle cohesion and rotational resistance. Fine particles with diameters of 0.5 mm, measured from the µCT images of the soil, were used in the simulation. The JKR adhesive model was incorporated to represent the cohesive behaviour and aggregation of soil particles (Table S2), thereby approximating the formation of soil clumps observed experimentally ^53–55^. A series of sensitivity tests was performed to assess the effects of static and dynamic friction, coefficient of restitution, surface energy, and rolling resistance. These parameters were systematically varied to calibrate the DEM-predicted forces against those measured in the *in vitro* experiments, and then used as input parameters for the *in vivo* DEM. Details were recorded in Supplementary Figure S4 – S9 and Table S3 – S8.

The geometry and boundary conditions of DEM simulations were configured to replicate the corresponding experimental setups in Section 4.2 and 4.3. For the *in vitro* DEM simulation, the skull was translated at a constant velocity of 0.01 m/s for 5 s to replicate the experimental burrowing test. For the DEM simulation used to mimic *in vivo* experiments, the same soil properties and contact parameters calibrated in the *in vitro* experiments were set as the input, while burrowing motion was derived from the corresponding *in vivo* force measurements. To approximate the extreme loading condition of full-body burrowing, the DEM-derived contact nodal forces were uniformly scaled using the ratio between the peak force measured *in vivo* and the corresponding one predicted by the DEM in three directions,

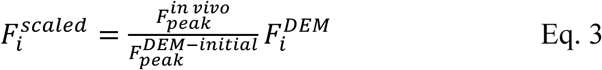

where *Fi^DEM^* is the force vector at node *i*, *Fpeak^in^ ^vivo^* is the peak resultant force measured during full-body burrowing at each direction, and *F^DEM^*^-*initial*^ is the corresponding peak resultant force predicted by the DEM at the initial stage. This procedure preserved the spatial distribution and relative magnitudes of the contact forces predicted by the DEM while matching the experimentally observed overall loading magnitude. The local forces were determined by summing the particle contacted nodal forces within regions corresponding to the sensor locations.

### 4.5 Discrete Element – Finite Element Coupling

The particle contact forces predicted by the DEM simulations were transferred to a finite element model of the skull to evaluate the resulting stress and strain distributions. The skull model was generated and meshed in Simpleware (X, Synopsys Inc., CA, US), with 8,241,581 tetrahedral solid elements, and imported into Mechanical APDL (2025 R2, ANSYS Inc., PA, US) for simulation. The material properties of the skull model were recorded in Table 1. Young’s moduli have not been experimentally determined for the skin or skeletal muscle of amphisbaenians. Therefore, material properties were approximated using experimentally measured values from comparable vertebrate tissues. At each DEM output timestep, the contact forces acting on the skull surface (skin) were extracted and mapped onto the corresponding nodes of the FE mesh. The neck region of the modelled skull was constrained to represent the experimental fixation conditions. A quasi-static structural analysis was then performed to calculate the von Mises stress and principal strain distributions within the skull.

### 4.6 Statistical Analysis

Agreement between the experimental and DEM-predicted force responses was quantified using the coefficient of determination (R^2^, Eq. 4) and the root mean square error (RMSE, Eq. 5) ^56^. The R^2^ value was used to assess the similarity in the overall force trends between the experimental and simulated data, while RMSE was used to quantify the average magnitude of the prediction error. Higher R^2^ values and lower RMSE values indicate better agreement between the DEM simulations and experimental measurements. Pearson’s correlation coefficient (Pearson r) was then used to quantify the strength and direction of the linear association between the experimental and simulated force profiles.

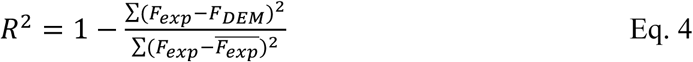

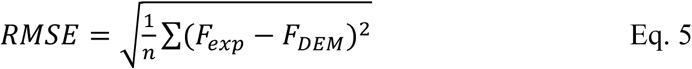

## Data availability

The authors declare that all data supporting the findings of this study are available within the manuscript and its supplementary materials.

## Acknowledgement

The authors wish to thank Dr Yulu Jiang (Mechanical Engineering, University College London, UK) for providing access to the high-performance workstation used for the computational simulations, and Ray Hong (Mechanical Engineering, University College London, UK) for assistance with setting up mechanical testing. The authors also wish to thank Sienna Griffin-Shaw (Bartlett Faculty of the Built Environment, University College London, UK) for 3D printing the skull models.

## Authorship Contribution Statement

**Jingrui Hu:** Writing – review & editing, Writing – original draft, Visualization, Validation, Methodology, Investigation, Formal analysis, Data curation, Conceptualization. **Anastasiia Maliuk:** Writing – review & editing, Data curation. **Ce Liang:** Writing – review & editing, Visualization, Data curation. **Sina Najjari:** Writing – review & editing, Methodology, Data curation. **Liwei Guo:** Writing – review & editing. **Sara-Adela Abad Guaman:** Writing – review & editing. **Dominique Adriaens:** Writing – review & editing, Funding acquisition. **Tiana Kohlsdorf:** Writing – review & editing, Funding acquisition. **Henrik Birkedal:** Writing – review & editing, Funding acquisition. **Anthony Herrel:** Writing – review & editing, Methodology, Formal analysis, Data curation, Funding acquisition. **Susan E. Evans:** Writing – review & editing, Funding acquisition, Conceptualization. **Mehran Moazen:** Writing – review & editing, Resources, Funding acquisition, Conceptualization.

