## Supplementary Information for "A Novel Computational Framework to Investigate the Biomechanics of Burrowing"

**Table S1.** Coefficient of determination (R^2^), Pearson correlation test (Pearson r) and root mean square error (RMSE) quantifying the differences in the DEM-predicted reaction force and nodal force responses with in vitro measured force.

| **Resultant Force** | | | |
| --- | --- | --- | --- |
| **Material Name** | ***R^2^*** | **Pearson *r*** | **RMSE (N)** |
| **PMMA** | 0.93 | 0.98 | 0.82 |
| **Sand** | 0.98 | 0.99 | 15.51 |
| **Soil** | 0.93 | 0.97 | 4.27 |
| **Nodal Force** | | | |
| **Material Name** | ***R^2^*** | **Pearson *r*** | **RMSE (N)** |
| **PMMA** | 0.54 | 0.87 | 0.12 |
| **Sand** | 0.81 | 0.92 | 4.18 |
| **Soil** | 0.92 | 0.99 | 0.30 |

**Table S2.** Contact interaction parameters used in DEM simulations.

| **Parameter** | **PMMA-PMMA** | **PMMA-Skull** |  | **Sand-Sand** | **Sand-Skull** |  | **Soil-Soil** | **Soil-Skull** |
| --- | --- | --- | --- | --- | --- | --- | --- | --- |
| **Static Friction** | 0.30 | 0.30 |  | 0.57 | 0.40 |  | 0.60 | 0.40 |
| **Dynamic Friction** | 0.30 | 0.30 |  | 0.30 | 0.30 |  | 0.30 | 0.40 |
| **Tangential Stiffness Ratio** | 1.00 | 1.00 |  | 1.00 | 1.00 |  | 1.00 | 1.00 |
| **Restitution** | 0.30 | 0.30 |  | 0.30 | 0.30 |  | 0.60 | 0.30 |
| **Surface energy (J/m⁻²)** | –– | –– |  | –– | –– |  | 0.04 | 0.02 |
| **Rolling friction** | 0.00 | –– |  | 0.20 | –– |  | 0.05 | –– |
| **Reference** | 1 |  |  | 2,3 |  |  | 4-6 |  |

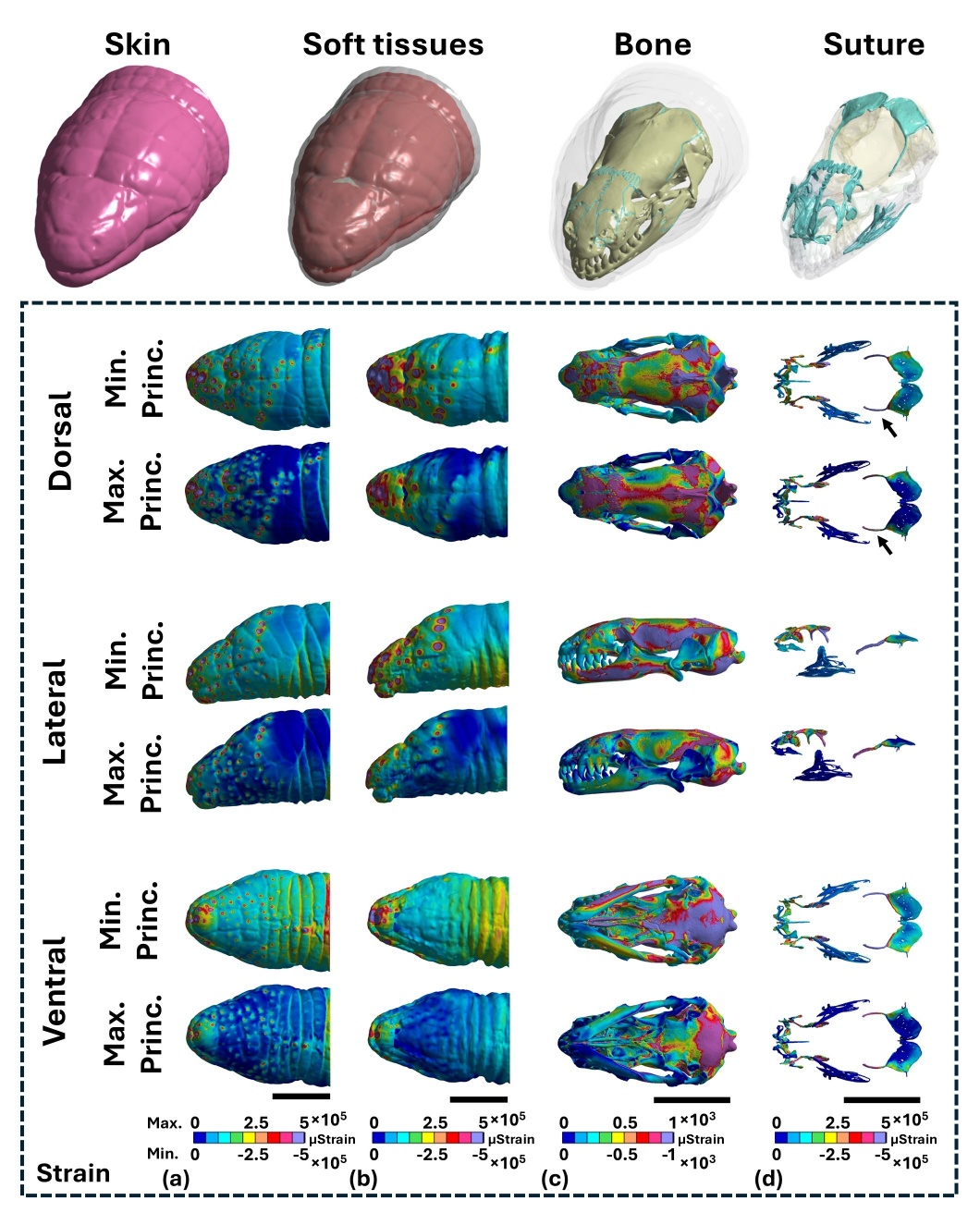

**Figure S1.** (a – d) Minimum and maximum principal strain distributions shown in dorsal, lateral, and ventral views. Scale bar = 3 mm.

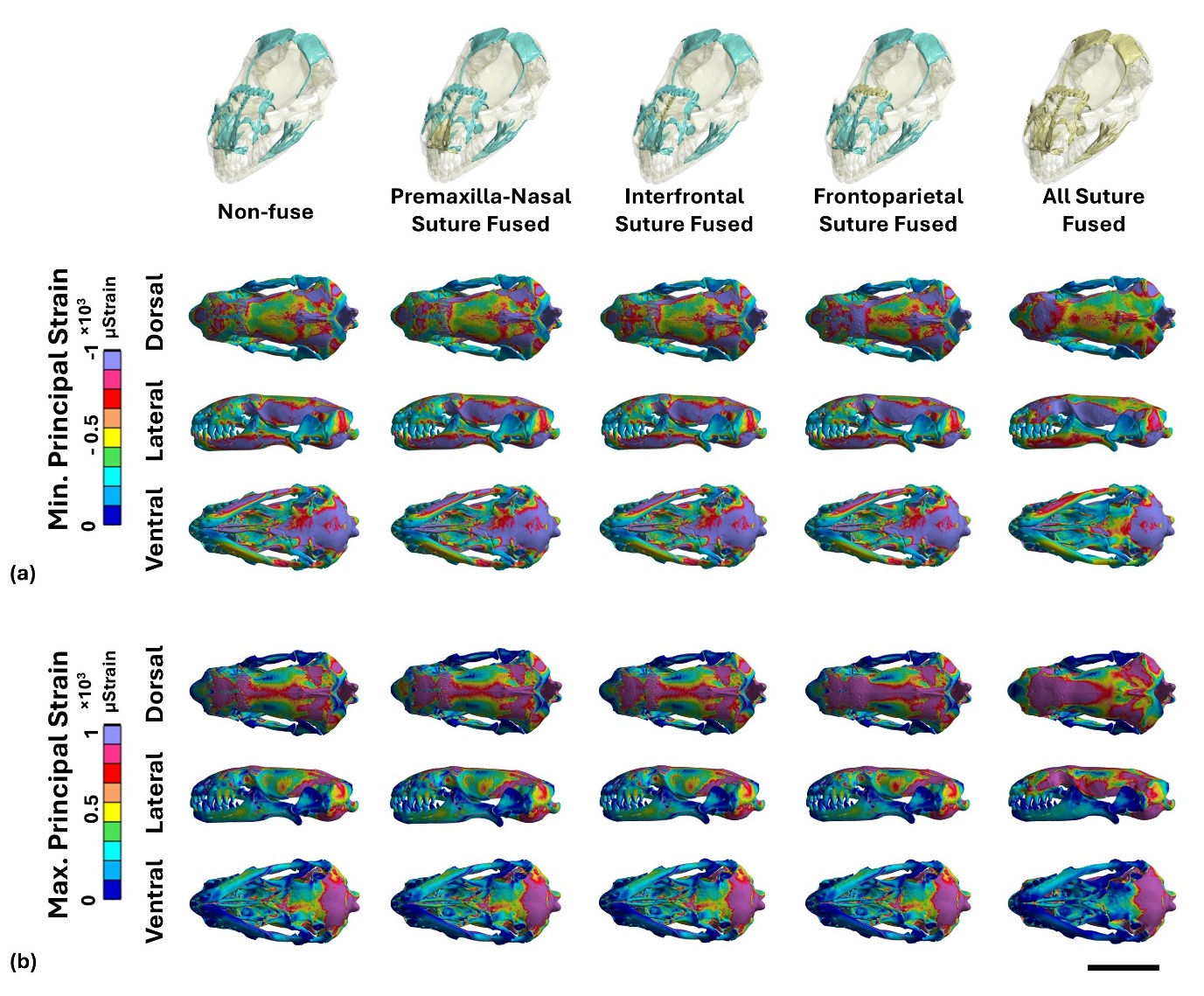

**Figure S2.** Minimum (a) and maximum (b) principal strain distribution of skull with different cranial suture fused conditions shown in dorsal, lateral, and ventral views. Scale bar = 3 mm.

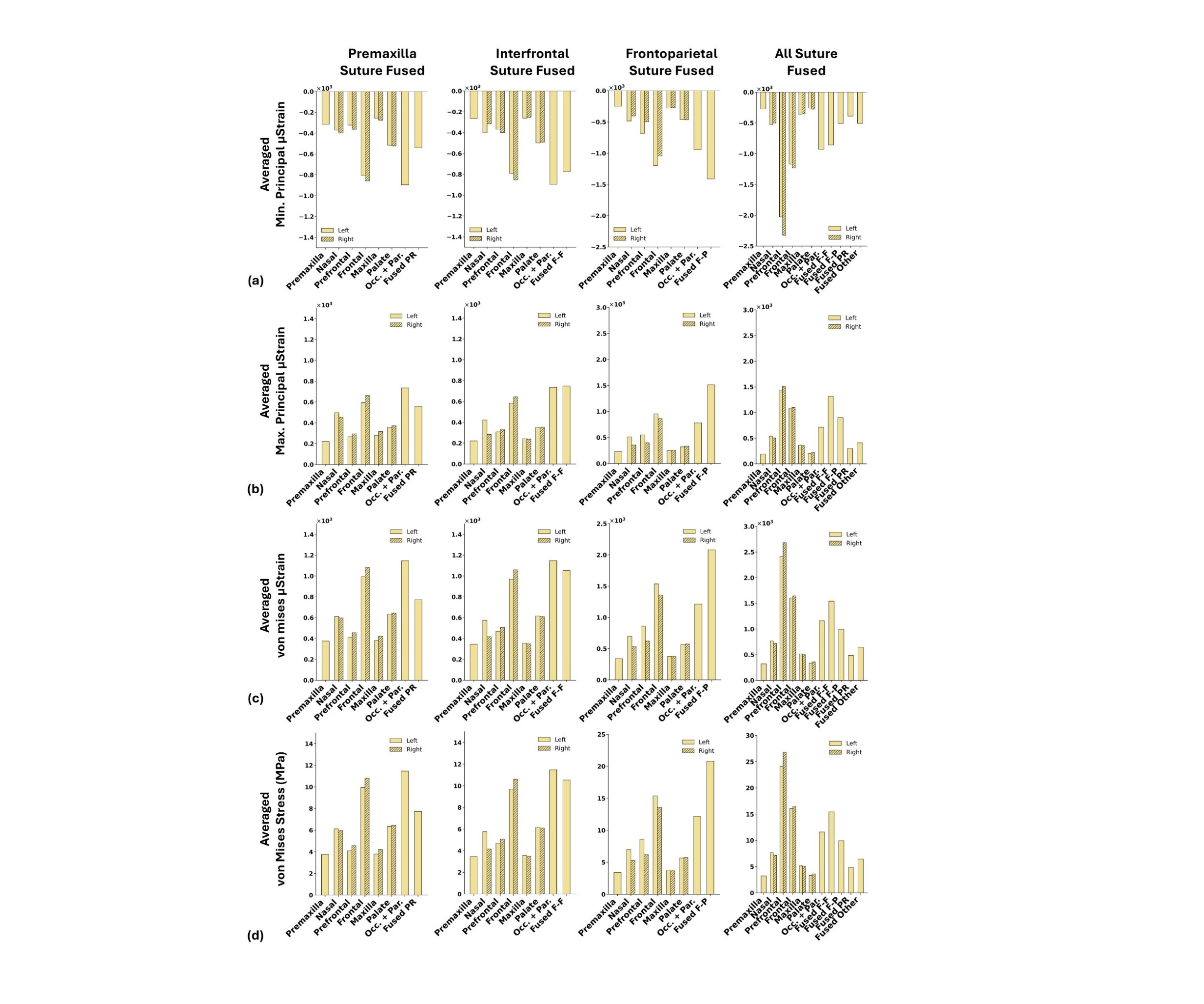

**Figure S3.** Average minimum and maximum principal strains and von Mises stress across cranial bone compartments at different suture fusion conditions. Pr. suture – premaxilla-nasal suture; F-F. suture – interfrontal suture; F-P suture – Frontoparietal suture.

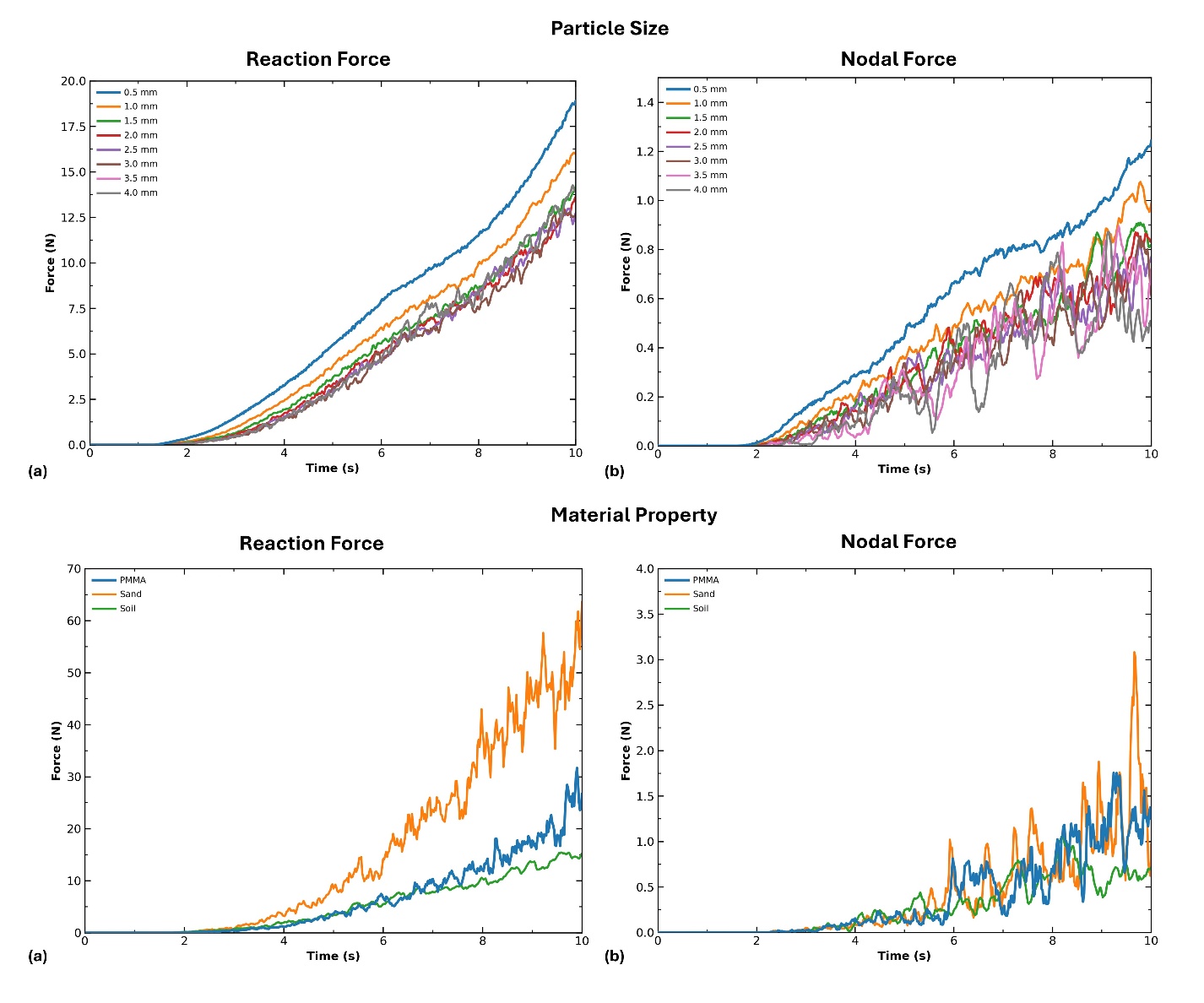

**Figure S4.** Sensitivity analysis of particle size in the in vitro DEM validation model. (a) Comparison of the reaction forces and (b) local nodal forces predicted, using different particle sizes.

**Table S3.** Coefficient of determination (R^2^) and root mean square error (RMSE) quantifying the differences in the DEM-predicted reaction force and nodal force responses with different particle sizes. 0.5 mm particle size was used as reference.

| **Reaction Force** | | | |
| --- | --- | --- | --- |
| **Reference label** | **Comparison label** | ***R^2^*** | **RMSE (N)** |
| 0.5 mm | 1.0 mm | 0.9422 | 1.3362 |
|  | 1.5 mm | 0.8410 | 2.2174 |
|  | 2.0 mm | 0.7867 | 2.5682 |
|  | 2.5 mm | 0.7732 | 2.6477 |
|  | 3.0 mm | 0.7396 | 2.8375 |
|  | 3.5 mm | 0.8124 | 2.4081 |
|  | 4.0 mm | 0.8124 | 2.4081 |
| **Nodal Force** | | | |
| **Reference label** | **Comparison label** | ***R^2^*** | **RMSE (N)** |
| 0.5mm | 1.0 mm | 0.9078 | 0.1172 |
|  | 1.5 mm | 0.7206 | 0.2040 |
|  | 2.0 mm | 0.6776 | 0.2191 |
|  | 2.5 mm | 0.5867 | 0.2481 |
|  | 3.0 mm | 0.5422 | 0.2611 |
|  | 3.5 mm | 0.4786 | 0.2786 |
|  | 4.0 mm | 0.3787 | 0.3042 |

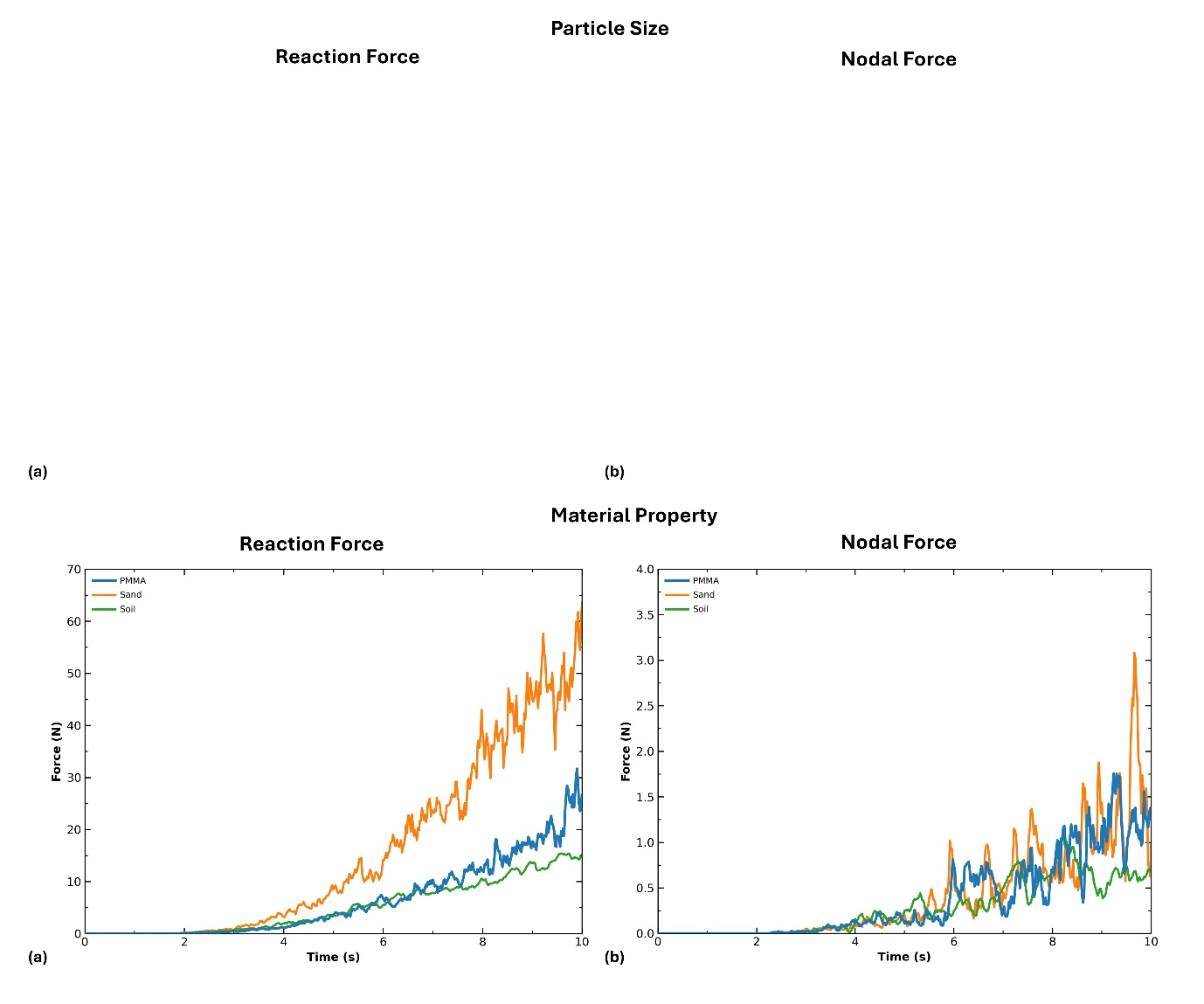

**Figure S5.** Comparison of the DEM-predicted reaction force (a) and nodal force (b) for three different material properties.

**Table S4.** Coefficient of determination (R^2^) and root mean square error (RMSE) quantifying the differences in the DEM-predicted reaction force and nodal force responses between PMMA and the sand and soil material models. PMMA was used as the reference, and 4 mm particle size was used for this convergence test.

| **Reaction Force** | | | |
| --- | --- | --- | --- |
| **Reference label** | **Comparison label** | ***R^2^*** | **RMSE (N)** |
| PMMA | Sand | -3.2426 | 9.3176 |
|  | Soil | 0.9008 | 1.4245 |
| **Nodal Force** | | | |
| **Reference label** | **Comparison label** | ***R^2^*** | **RMSE (N)** |
| PMMA | Sand | 0.5434 | 0.2980 |
|  | Soil | 0.6036 | 0.2776 |

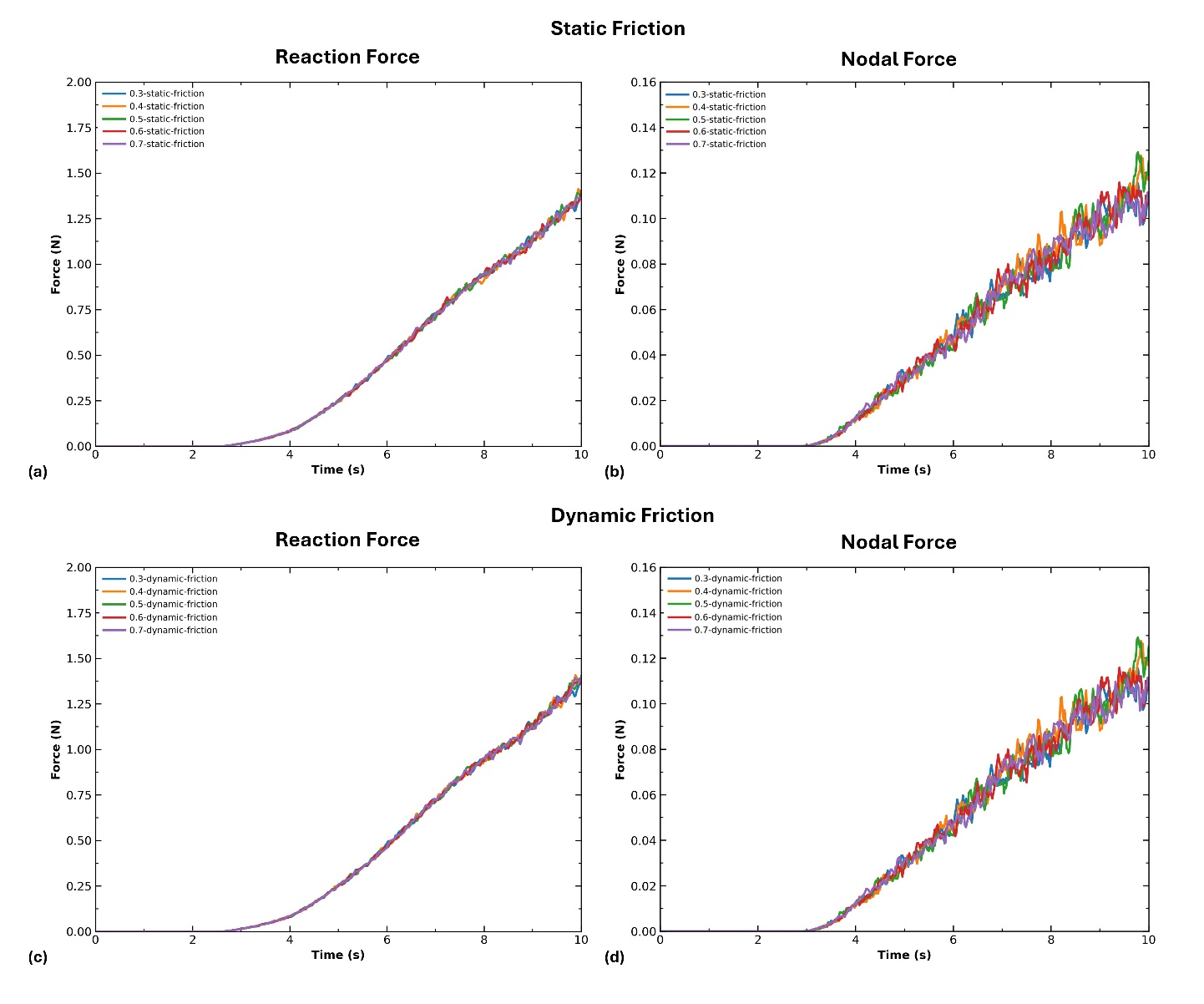

**Figure S6.** Comparison of the DEM-predicted reaction force and nodal force to the static and dynamic friction coefficients. Static friction coefficients ranging from 0.3 to 0.7 are shown in (a) and (b), while dynamic friction coefficients ranging from 0.3 to 0.7 are shown in (c) and (d). Reaction force responses are presented in (a) and (c), and nodal force responses in (b) and (d). 1.5 mm soil particle was used for this convergence test.

**Table S5.** Coefficient of determination (R^2^) and root mean square error (RMSE) comparing the DEM-predicted reaction force and nodal force responses for different static and dynamic friction coefficients. The models with a friction coefficient of 0.3 were used as the reference.

| **Reaction Force** | | | |
| --- | --- | --- | --- |
| **Reference label** | **Comparison label** | ***R^2^*** | **RMSE (N)** |
| 0.3-static-friction | 0.4-static-friction | 0.9991 | 0.0134 |
|  | 0.5-static-friction | 0.9992 | 0.013 |
|  | 0.6-static-friction | 0.9994 | 0.0114 |
|  | 0.7-static-friction | 0.9993 | 0.0117 |
| **Reference label** | **Comparison label** | ***R^2^*** | **RMSE (N)** |
| 0.3-dynamic-friction | 0.4-dynamic-friction | 0.9989 | 0.0148 |
|  | 0.5-dynamic-friction | 0.9993 | 0.0119 |
|  | 0.6-dynamic-friction | 0.9992 | 0.0124 |
|  | 0.7-dynamic-friction | 0.9991 | 0.0136 |
| **Nodal Force** | | | |
| **Reference label** | **Comparison label** | ***R^2^*** | **RMSE (N)** |
| 0.3-static-friction | 0.4-static-friction | 0.9846 | 0.0048 |
|  | 0.5-static-friction | 0.9886 | 0.0042 |
|  | 0.6-static-friction | 0.9875 | 0.0044 |
|  | 0.7-static-friction | 0.9873 | 0.0044 |
| **Reference label** | **Comparison label** | ***R^2^*** | **RMSE (N)** |
| 0.3-dynamic-friction | 0.4-dynamic-friction | 0.9835 | 0.005 |
|  | 0.5-dynamic-friction | 0.9849 | 0.0048 |
|  | 0.6-dynamic-friction | 0.9907 | 0.0038 |
|  | 0.7-dynamic-friction | 0.9881 | 0.0042 |

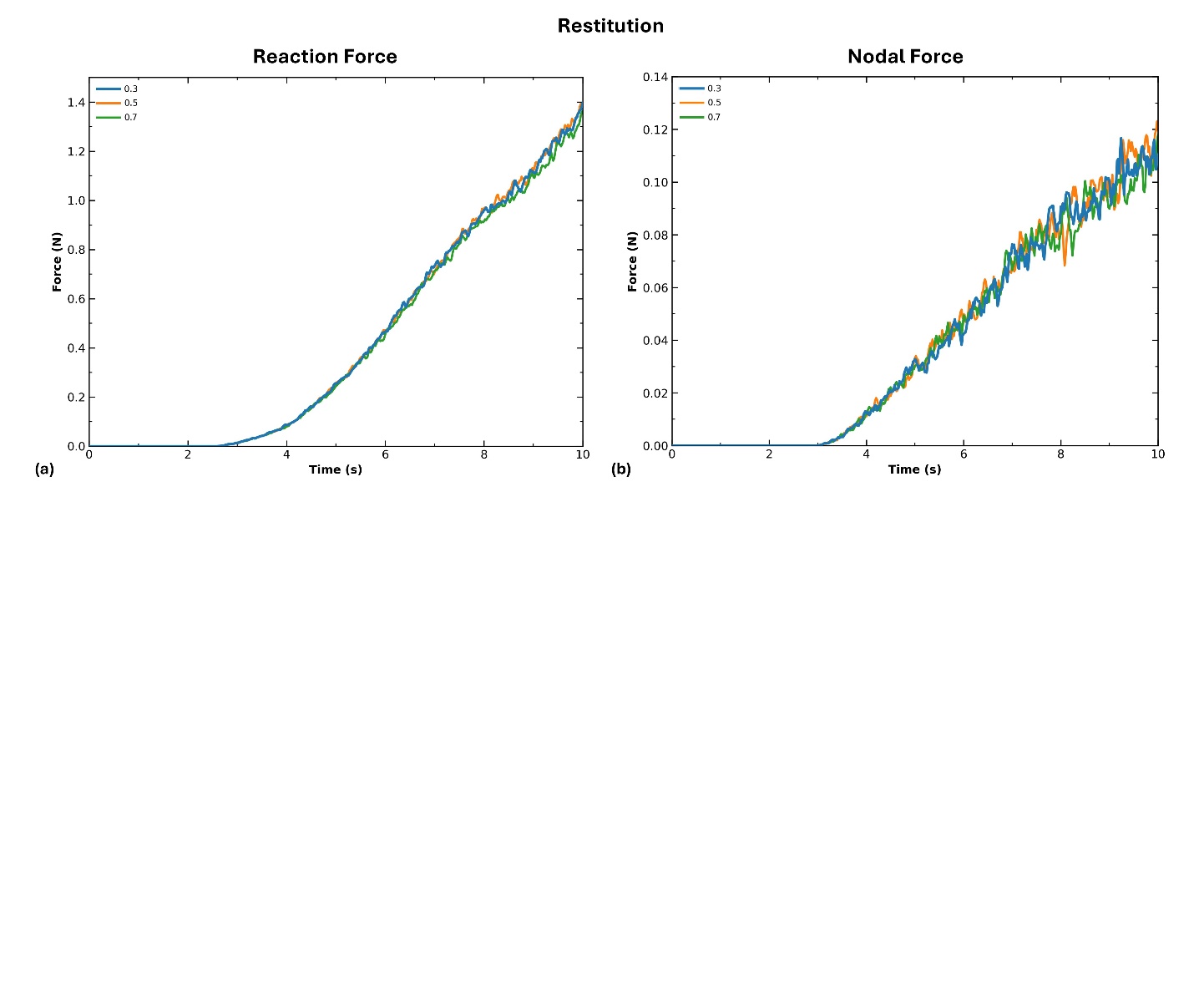

**Figure S7.** Comparison of the DEM-predicted reaction force and nodal force to restitution ranging from 0.3 to 0.7 are shown in (a) and (b).

**Table S6.** Coefficient of determination (R^2^) and root mean square error (RMSE) comparing the DEM-predicted reaction force and nodal force responses for restitution. The models with restitution of 0.3 were used as the reference.

| **Reaction Force** | | | |
| --- | --- | --- | --- |
| **Reference label** | **Comparison label** | ***R^2^*** | **RMSE (N)** |
| 0.3 | 0.5 | 0.9994 | 0.0113 |
|  | 0.7 | 0.9981 | 0.0196 |
| **Nodal Force** | | | |
| **Reference label** | **Comparison label** | ***R^2^*** | **RMSE (N)** |
| 0.3 | 0.5 | 0.9857 | 0.0047 |
|  | 0.7 | 0.9885 | 0.0042 |

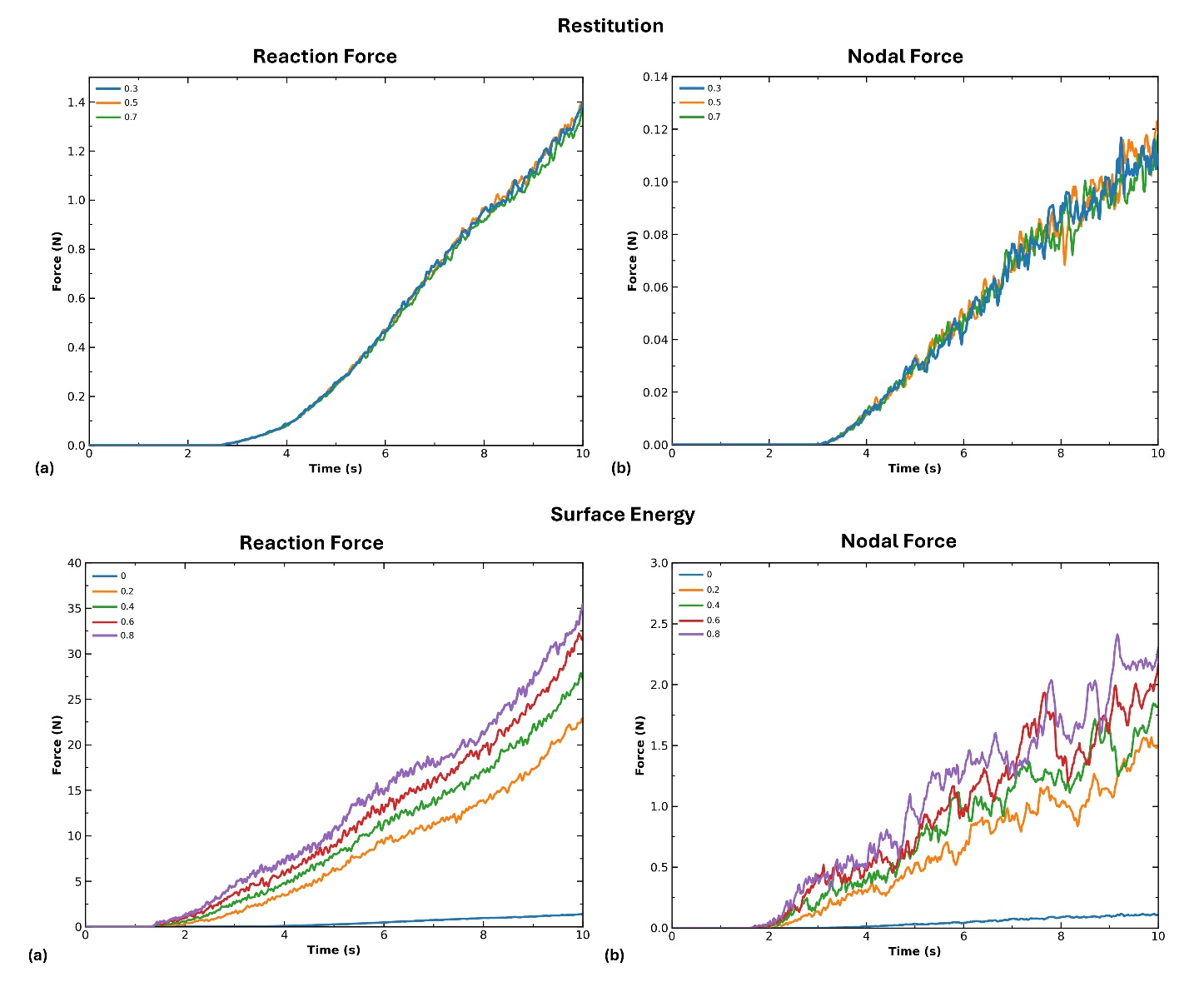

**Figure S8.** Comparison of the DEM-predicted (a) reaction force and (b) nodal force for surface energies ranging from 0 to 0.8 J/m^2^, at intervals of 0.2 J/m^2^.

**Table S7.** Coefficient of determination (R^2^) and root mean square error (RMSE) comparing the DEM-predicted reaction force and nodal force responses for surface energies ranging from 0 to 0.8 J/m^2^, at intervals of 0.2 J/m^2^. The models with restitution of 0.8 J/m^2^ were used as the reference.

| **Reaction Force** | | | |
| --- | --- | --- | --- |
| **Reference label** | **Comparison label** | ***R^2^*** | **RMSE (N)** |
| 0.8 | 0 | -1.2974 | 15.2724 |
|  | 0.2 | 0.6563 | 5.9074 |
|  | 0.4 | 0.8703 | 3.6291 |
|  | 0.6 | 0.9693 | 1.7662 |
| **Nodal Force** | | | |
| **Reference label** | **Comparison label** | ***R^2^*** | **RMSE (N)** |
| 0.8 | 0 | -1.4177 | 1.1561 |
|  | 0.2 | 0.5517 | 0.4978 |
|  | 0.4 | 0.7918 | 0.3393 |
|  | 0.6 | 0.9206 | 0.2095 |

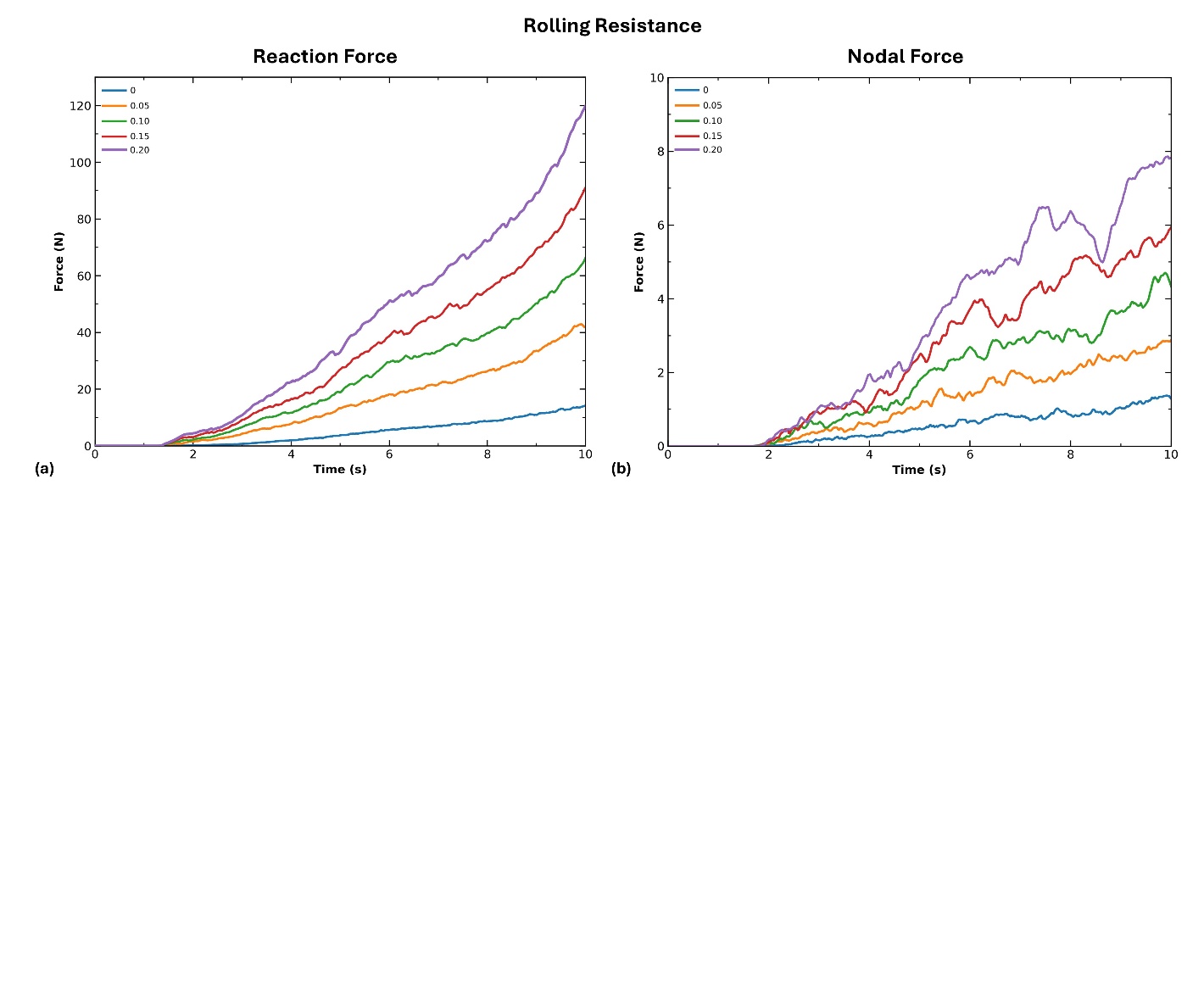

**Figure S9.** Comparison of the DEM-predicted reaction force and nodal force to rolling resistance ranging from 0 to 0.2, with intervals of 0.05, are shown in (a) and (b).

**Table S8.** Coefficient of determination (R^2^) and root mean square error (RMSE) comparing the DEM-predicted reaction force and nodal force responses for rolling resistance. The models with rolling resistance of 0.20 were used as the reference.

| **Reaction Force** | | | |
| --- | --- | --- | --- |
| **Reference label** | **Comparison label** | ***R^2^*** | **RMSE (N)** |
| 0.20 | 0.00 | -0.8526 | 46.3704 |
|  | 0.05 | 0.0395 | 33.3895 |
|  | 0.10 | 0.5291 | 23.3786 |
|  | 0.15 | 0.8636 | 12.5808 |
| **Nodal Force** | | | |
| **Reference label** | **Comparison label** | ***R^2^*** | **RMSE (N)** |
| 0.20 | 0.00 | -0.7537 | 3.4891 |
|  | 0.05 | -0.0372 | 2.6833 |
|  | 0.10 | 0.4817 | 1.8968 |
|  | 0.15 | 0.8448 | 1.0379 |

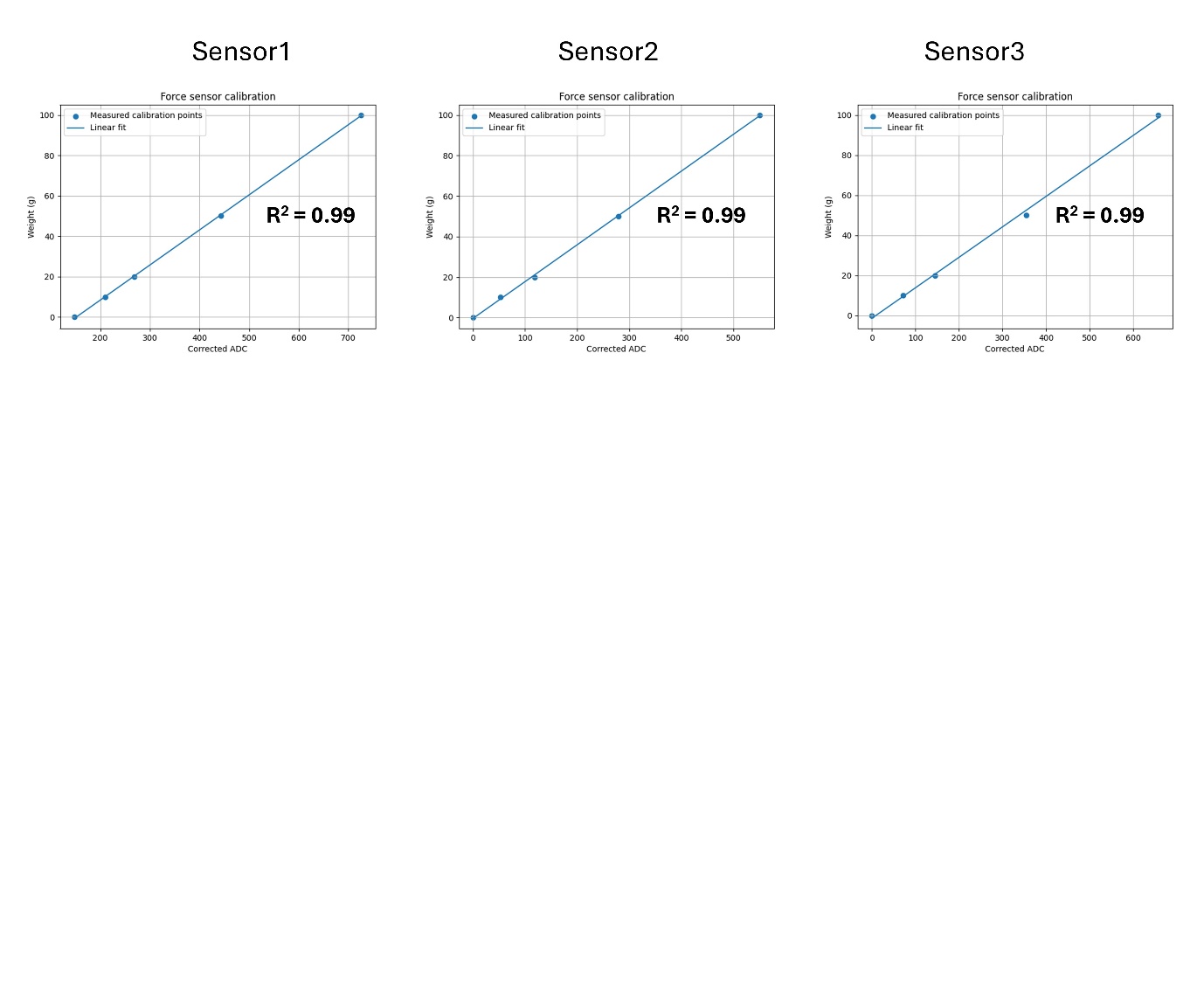

**Figure S10.** Calibration of the force sensor across three repeated trials, showing the linear relationship between applied weight and ADC values (R² = 0.99).
